# The human frontal operculum is involved in visuomotor performance monitoring

**DOI:** 10.1101/2021.10.21.465314

**Authors:** Felix Quirmbach, Jakub Limanowski

**Affiliations:** Faculty of Psychology, Technische Universität Dresden, Dresden, Germany; Centre for Tactile Internet with Human-in-the-Loop, Technische Universität Dresden, Dresden, Germany

**Keywords:** Action, frontal operculum, sensorimotor integration, performance monitoring, visuo-proprioceptive integration

## Abstract

For adaptive goal-directed action, the brain needs to monitor action performance and detect errors. The corresponding information may be conveyed via different sensory modalities; for instance, visual and proprioceptive body position cues may inform about current manual action performance. Thereby, contextual factors such as the current task set may also determine the relative importance of each sensory modality for action guidance. Here, we analysed human behavioral, fMRI, and MEG data from two VR-based hand-target phase matching studies to identify the neuronal correlates of performance monitoring and error processing under instructed visual or proprioceptive task sets. Our main result was a general, modality-independent response of the bilateral FO to poor phase matching accuracy, as evident from increased BOLD signal and increased gamma power. Furthermore, functional connectivity of the bilateral FO to the right PPC increased under a visual vs proprioceptive task set. These findings suggest that the bilateral FO generally monitors manual action performance; and, moreover, that when visual action feedback is used to guide action, the FO may signal an increased need for control to visuomotor regions in the right PPC following errors.

## Introduction

To effectively perform goal-directed action in the environment, the brain needs to monitor motor performance and detect errors, so that it can enable adaptive changes in behavior (Diedrichsen, 2005; Klein et al., 2007; Suminski et al., 2007; Ullsperger et al., 2014). During performance monitoring, the predicted outcome of one’s actions selected based on current goals (i.e., task set) is compared with actual sensory feedback, and behavioural changes are initiated if a mismatch between both is detected (Ullsperger et al., 2014). The neurofunctional basis of performance monitoring and error correction has been illuminated by recent brain imaging and electrophysiological work. Specifically, a ‘salience network’ comprising, among others, the dorsal anterior cingulate cortex, the bilateral insular cortex and the inferior frontal gyri, is assumed to integrate sensory input, responding to behaviourally salient stimuli—behavioural errors—with increased activation (Ham et al., 2013; Seeley et al., 2007; Sridharan et al., 2008; Uddin, 2021). Thereby regions like e.g. the frontal operculum (FO, also anterior insular cortex, IC(Billeke et al., 2020; Cieslik et al., 2015; Higo et al., 2011; Klein et al., 2013; Sridharan et al., 2008) may signal a need for increased cognitive control to the executive control network, consisting (among other regions) of the lateral prefrontal cortices, the posterior parietal cortex, pre-supplementary motor area and the inferior parietal lobule(Uddin, 2021; Ullsperger et al., 2010). This network, in turn, may direct attentional resources to the relevant stimuli, driving behavioural adaptions (Menon & Uddin, 2010; Sridharan et al., 2008).

Notably, action performance and error may be conveyed via different sensory modalities; in manual action, for instance, via visual and proprioceptive cues about body position. In the context of body representation for action, visual and proprioceptive body position cues can be weighted depending on the current context; e.g., based on their relative relevance for the specific task at hand (Lebar et al., 2017; Sober & Sabes, 2005; van Beers et al., 1999). Recently, we have used virtual reality (VR) to examine this contextual sensory weighting during action under conflicting visual (virtual) and proprioceptive (real, unseen) body position feedback. Our functional magnetic resonance imaging (fMRI) and magnetoencephalography (MEG) studies (Limanowski et al., 2020; Limanowski & Friston, 2020) specifically shed light on the effects of adopting a visual vs proprioceptive attentional set during goal-directed manual action tasks, demonstrating that participants’ can prioritise either modality over the other; and we observed corresponding changes of neuronal gain in the respective sensory (visual and proprioceptive) brain regions. However, while the effects of adopting an attentional set on sensory processing could be seen clearly, these studies did not investigate the specific neural correlates of flexible performance monitoring in these settings.

Here, we aimed to close this gap. We therefore re-analysed the behavioural, fMRI, and MEG data from the above studies, correlating participants’ task performance with hemodynamic and oscillatory activity. Based on the above literature, we expected task inaccuracy to be reflected by activity in the performance monitoring network and, potentially, also in fronto-parietal attentional areas. Our second research question was whether performance monitoring would be modality specific (i.e., involve different brain regions when vision vs proprioception was task relevant) or general. Therefore, we also tested for task set dependent differences in brain activity and connectivity.

## Materials and Methods

### Participants

For this study, we reanalysed fMRI and MEG data acquired by (Limanowski et al., 2020; Limanowski & Friston, 2020). Healthy, right-handed volunteers with normal or corrected-to-normal vision participated in both experiments after providing written informed consent. The fMRI study included 16 subjects (8 female, mean age 27, range 21-37), the MEG study in included 18 subjects (9 female, mean age 29, range 21-39). Both experiments were approved by the local research ethics committee (University College London) and conducted in accordance with these approvals.

### Experimental design and task

Participants wore an MR compatible data glove (5DT Data Glove MRI, 1 sensor per finger, 8 bit flexure resolution per sensor, 60 Hz sampling rate, communication with the PC via USB) on their right hand. The glove measured each finger’s flexion via sewn-in optical fibre cables, and was carefully calibrated to fit each participant’s movement range prior to scanning. Recorded hand movement data was used to control a photorealistic virtual hand (VH) model, moving in accordance to the participant’s hand movements and presented as part of a virtual reality task environment. This virtual environment, consisting of the VH, a fixation dot and task instructions, was created in the open-source 3D computer graphics software Blender (http://blender.org). The environment was presented via a projector on a screen (for details see (Limanowski et al., 2020; Limanowski & Friston, 2020).

Participants were instructed to perform repetitive right-hand grasping movements, paced by oscillatory (0.5 Hz) size changes (12%) of the central fixation dot, resulting in a non-spatial phase matching task: Thus, participants had to match the fully open hand position with the biggest dot size and, conversely, the fully closed hand with minimal dot size. They performed the task in movement blocks of 32 s (16 close-and-open movements; the last movement was signaled by brief blinking of fixation dot), separated by 16 s rest periods during which only the fixation dot was visible. All participants trained extensively before scanning. Note that this task was, therefore, not designed to investigate visuomotor adaptation or learning, but maintaining hand-target phase matching during a sustained visual vs proprioceptive attentional task set.

Before the start of each movement block, participants were instructed to match the phase of the fixation dot with either the seen VH model or their unseen real hand. In half of the conditions, a lag was introduced to the virtual hand’s movements; i.e., the virtual hand movements lagged behind the actually executed movements. In the fMRI experiment, a lag of 267 ms was introduced in the second half of each movement block; in the MEG experiment, a lag of 500 ms was presented as a separate block. Note that under incongruence, only one modality could be aligned with the target phase, which resulted in a misalignment of the other one. The task instruction (‘VIRTUAL’ or ‘REAL’) were presented 2.5 s before start of each movement block for 2 s, and the color of the fixation dot reminded participants of the current condition thorough the block. In the fMRI experiment (Limanowski & Friston, 2020), we had additionally varied the visibility (high or low) of the virtual hand during half of the movement blocks. However, we found no differences in performance between different visibility levels; and in our present reanalysis, there were no significant differences between visibility levels either. Therefore, we present the differential fMRI task contrasts in terms of VH vs RH task, summing over high and low visibility levels in each condition. In sum, despite minor technical differences between fMRI and MEG experiments, both can be described as a balanced 2 × 2 factorial design with the factors *task* (VH vs RH) and *congruence* (congruent vs incongruent).

### Behavioural Data Analysis

In our previous analyses, we examined the neuronal correlates of the instructed task set; and only analysed condition specific differences in average performance (Limanowski et al., 2020; Limanowski & Friston, 2020). In the present study, we examined the neuronal correlates of phase matching *accuracy* (i.e., *fluctuations* around those average performances).

To quantify hand-target phase matching (in)accuracy, we calculated the root mean square error (RMSE) of the difference between the target position (i.e., the position within the oscillatory cycle) and the position of the task relevant hand (i.e., the position within the grasping cycle, averaging the recorded finger position data per hand). Thus, the virtual hand position was evaluated for the VH condition movements and the real hand position for the RH condition. For construction of the fMRI/MEG regressors, we binned the resulting RSME values into 1 s time windows, each centred on a time point of minimum or maximum target size, corresponding to the hand fully closed or opened if moved synchronously with the target. To focus on within-subject fluctuations in performance, rather than between-subject differences, the overall RSME across the entire experiment was normalized for each single subject (i.e. minimum and maximum performance error value was equal across participants; 0 and 1, respectively). The resulting RSME values were assigned to one regressor per experimental condition (VH congruent, VH incongruent, RH congruent, RH incongruent), and demeaned separately to reflect only variation around the condition mean. To evaluate if the variance of the phase matching differed between the VH and RH conditions, we calculated a paired t-test on the participants’ variance of phase matching RMSE within each condition.

The amplitude of the hand movement at each time point was calculated via a cubic spine interpolation of the respective minimum and maximum hand position values in each time window. The resulting time series was de-meaned per condition as well, and used as noise regressor for the following fMRI and MEG analysis (see below). Additionally, we tested for correlations between performance error and movement amplitude by calculating the Pearson correlation coefficient of both regressors for each participant; and testing it for significance (i.e., significant difference from zero) with a t-test on the group level. Similarly, we calculated the correlation between performance error and fMRI head movements (realignment parameters), via subject-level Pearson correlation and group-level t-test, adjusted for multiple comparisons for the six realignment parameters. All analyses were performed using MATLAB (MathWorks, Natick, MA, United States).

### FMRI Data Preprocessing and Analysis

All analyses were performed using MATLAB (MathWorks, Natick, MA, United States) and SPM12.6 (Wellcome Trust Centre for Neuroimaging, University College London, https://www.fil.ion.ucl.ac.uk/spm/).

We reused the preprocessed fMRI data by (Limanowski & Friston, 2020). The fMRI data had been acquired using a 3T scanner (Magnetom TIM Trio, Siemens), equipped with a 64-channel head coil. T_2_*-weighted images were acquired using gradient echo-planar imaging sequence (voxel size= 3 × 3 × 3 mm^3^, matrix size= 64 × 72, TR = 3.36 s, TE = 30 ms, flip angle = 90°).

We fitted a general linear model (GLM, 128 s high-pass filter) to each participant. Each condition (VH, RH) was modelled with a boxcar function as a 32 s movement block; we added a parametric modulator (1/-1) to each condition encoding the first half of each block as congruent (−1) and the second half as incongruent (1) movement periods. Additionally, we included a regressor encoding the (de-meaned) RSME values for each conditions; the values were re-sampled to match the 3.36 s scan length prior to this. Regressors modelling the task instructions and movement amplitude were added to the GLM alongside the realignment parameters as regressors of no interest.

For each subject, we calculated contrast images of each RSME regressor against the baseline. These were then entered into a group-level flexible factorial design, with the factors *task* (VH or RH) and *congruence* (congruent, incongruent), and an additional factor modelling the subject constants. To assess potential differences between congruent and incongruent movement periods, we calculated separate first-level GLMs, in which the RSME values of the second movement half were inverted; this effectively encoded the contrast congruent-incongruent. The resulting contrast images were entered into an analogous group-level GLM as described above.

Group-level results were assessed for statistical significance using a voxel-wise threshold of *p* < 0.05, family-wise error (*p*_FWE_) corrected for multiple comparisons. We projected the resulting statistical maps onto the mean normalized structural image or rendered it on SPM12’s brain template. The unthresholded T-maps corresponding to the contrasts reported here can be inspected online at https://neurovault.org/collections/GGWQTGSI. For anatomical reference we used the SPM Anatomy toolbox (Eickhoff et al., 2005).

### MEG Data Preprocessing and Analysis

MEG signals had been acquired using a 275-channel whole-head setup with third-order gradiometers (CTF Omega, CTF MEG International Services LP, Coquitlam, Canada) at a sampling rate of 600 Hz. Following the original analysis by (Limanowski et al., 2020), the MEG data were high-pass filtered (1 Hz), downsampled to 300 Hz, and epoched into trials of 2 s each (each corresponding to a full target oscillation/grasping cycle).

In the main (sensor space) MEG data analysis, we looked for spectral power differences under ‘steady-state’ assumptions; i.e., treating the spectral profile as a ‘snapshot’ of performance dependent responses as manifest in quasi-stationary power spectra (Donner & Siegel, 2011; Friston et al., 2019; Moran et al., 2008). We computed trial-by-trial power spectra in the 0-98 Hz range using a multi-taper spectral decomposition (Thomson, 1982) with a spectral resolution of +-2 Hz. The spectra were log-transformed, converted to volumetric scalp × frequency images—one image per trial—with two spatial and one frequency dimension (Kilner & Friston, 2010), and smoothed with a Gaussian kernel with full width at half maximum of 8 mm × 8 mm × 4 Hz. The resulting images were entered into a general linear model (GLM) using a within-subject ANOVA with the respective RSME values as a covariate (first-level analysis). As in the fMRI analysis, movement amplitude was moreover included as a covariate of no interest to capture movement related fluctuations. Contrast images were then calculated for each condition’s accuracy covariate. These contrast images were then entered into a group-level GLM using a flexible factorial design including the two within-subject experimental factors (task and congruence), and a factor modelling the between subject variance. The statistical parametric maps obtained from the respective group-level contrasts were evaluated for significant effects using a threshold of *p* < 0.05, family-wise error (*p*_FWE_) corrected for multiple comparisons at the peak (voxel) level.

As a post-hoc analysis, source localization of trial-by-trial correlation of gamma band power with performance error was performed using a variational Bayesian approach with multiple sparse priors (Litvak & Friston, 2008). Source localization was performed in the 34-88 Hz range (which was the range of effects in the spectral analysis thresholded at *p* < 0.001, uncorrected). As we had already performed an analogous localization on the fMRI data (see above), we could use the superior spatial acuity of fMRI to improve MEG source localization; i.e., the fMRI activations (thresholded at *p* < 0.001, uncorrected) were used as empirical (spatial) priors for the Bayesian inversion routine (Henson et al., 2010; López et al., 2014). For comparison, we also reconstructed the sources using a Bayesian beamforming approach (Belardinelli et al., 2012). This produced very similar results; i.e., the strongest effects were localized to the bilateral inferior frontal gyri, including the FO (a further, weaker source was localized to the primary visual cortex). The results of this source localization were summarized as 3D images and entered into a group-level t-test. Since the significance of the effects on spectral responses had already been established with the sensor space analysis, the ensuing statistical parametric maps were displayed at a threshold of *p* < 0.05, uncorrected, rendered on SPM’s smoothed average brain template. The unthresholded T-map corresponding to the source localization can be inspected online at https://neurovault.org/collections/GGWQTGSI.

### FMRI functional connectivity analysis

In our main analysis (see above), we identified brain areas that showed a significance response to phase matching inaccuracy. The fMRI and MEG results consistently highlighted the bilateral FO (while further fMRI activations were found in the SMA and the dlPFC).

In our original analyses (Limanowski et al., 2020; Limanowski & Friston, 2020), the FO did not show any task specific effects (i.e., activity differences between VH and RH tasks) per se, neither did the SMA or the dlPFC. However, following the clear response of the FO to poor task performance in general, we now asked whether these areas would change their connectivity to other (potentially, task relevant) brain areas depending on whether the inaccuracy was registered during the VH or RH task; i.e., while participants focused either on visual (VH) or proprioceptive (RH) action feedback.

To answer this question, we used psychophysiological interaction (PPI) analysis for fMRI data. This analysis aims to explain neuronal activity other brain areas in terms of an interaction between psychological factors (the specific task condition) and physiological factors (the BOLD signal time course in the region of interest (Friston et al., 1997; O’Reilly et al., 2012). The resulting interaction (PPI) reveals voxels in the brain increase their connectivity with a specific seed region in a given context; e.g., in a specific task condition. Note that task dependent changes in connectivity per se (i.e., between VH and RH task sets) were identified in both fMRI and MEG data sets (Limanowski et al., 2020; Limanowski & Friston, 2020). However, in the fMRI data, the SPM approach allowed us to select a spatially isolated volume of interest (i.e., voxels from the FO) that was not part of the original connectivity analysis. This was not analogously possible for the MEG data, which in the original connectivity analysis were modelled on the whole-scalp level (Limanowski et al., 2020). Therefore, we limited the connectivity analysis to the fMRI data.

For the PPI analysis, we calculated separate GLMs with concatenated runs for each participant, and thus identified subject-specific peaks of the main effects observed on the group level. The individual peaks were defined as the maximum effect within a 10 mm radius sphere of the respective group-level maximum (see Table 1). From these individual peaks, we extracted the BOLD signal of the seed regions, as the first eigenvariate of activity across all voxels in a 4 mm radius sphere centred on the participant-specific peak. For three subjects where no effect could be identified for the specific SMA seed region as well as one case were no effect was found for the dlPFC region, we resorted to the group level maximum for seed region localization.

**Table 1.**
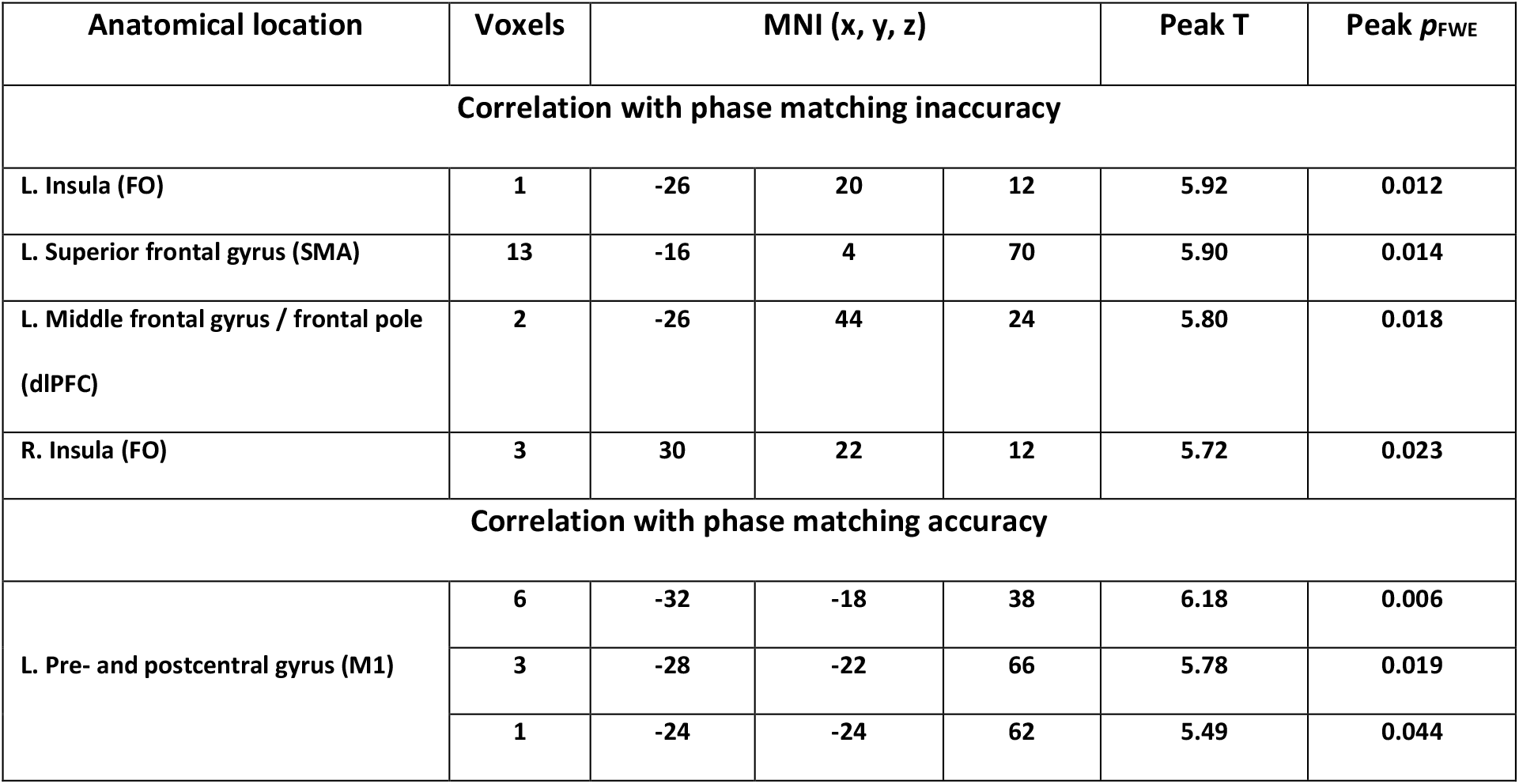
Significant (*p*_FWE_ < 0.05) activations for all reported fMRI contrasts.

| Anatomical location | Voxels | MNI (x, y, z) | | | Peak T | Peak $p_{FWE}$ |
| --- | --- | --- | --- | --- | --- | --- |
| Correlation with phase matching inaccuracy |  |  |  |  |  |  |
| L. Insula (FO) | 1 | -26 | 20 | 12 | 5.92 | 0.012 |
| L. Superior frontal gyrus (SMA) | 13 | -16 | 4 | 70 | 5.90 | 0.014 |
| L. Middle frontal gyrus / frontal pole<br>(dlPFC) | 2 | -26 | 44 | 24 | 5.80 | 0.018 |
| R. Insula (FO) | 3 | 30 | 22 | 12 | 5.72 | 0.023 |
| Correlation with phase matching accuracy |  |  |  |  |  |  |
| L. Pre- and postcentral gyrus (M1) | 6 | -32 | -18 | 38 | 6.18 | 0.006 |
|  | 3 | -28 | -22 | 66 | 5.78 | 0.019 |
|  | 1 | -24 | -24 | 62 | 5.49 | 0.044 |

The SPM12.6 PPI routine was then used to form the interaction between the psychological factor and the seed region’s summarized BOLD signal time course. Note that while the seed regions were identified per their significant response to phase matching inaccuracy (Fig. 2), our psychological factor was the task set; i.e., the instructed hand modality at the beginning of each movement block (VH vs RH; pooled over different levels of virtual hand visibility, see above). After forming the interaction term, a second GLM was constructed for each participant, including the interaction, the seed region’s extracted signal, the task set and the realignment parameters as regressors of no interest.

**Figure 1.**
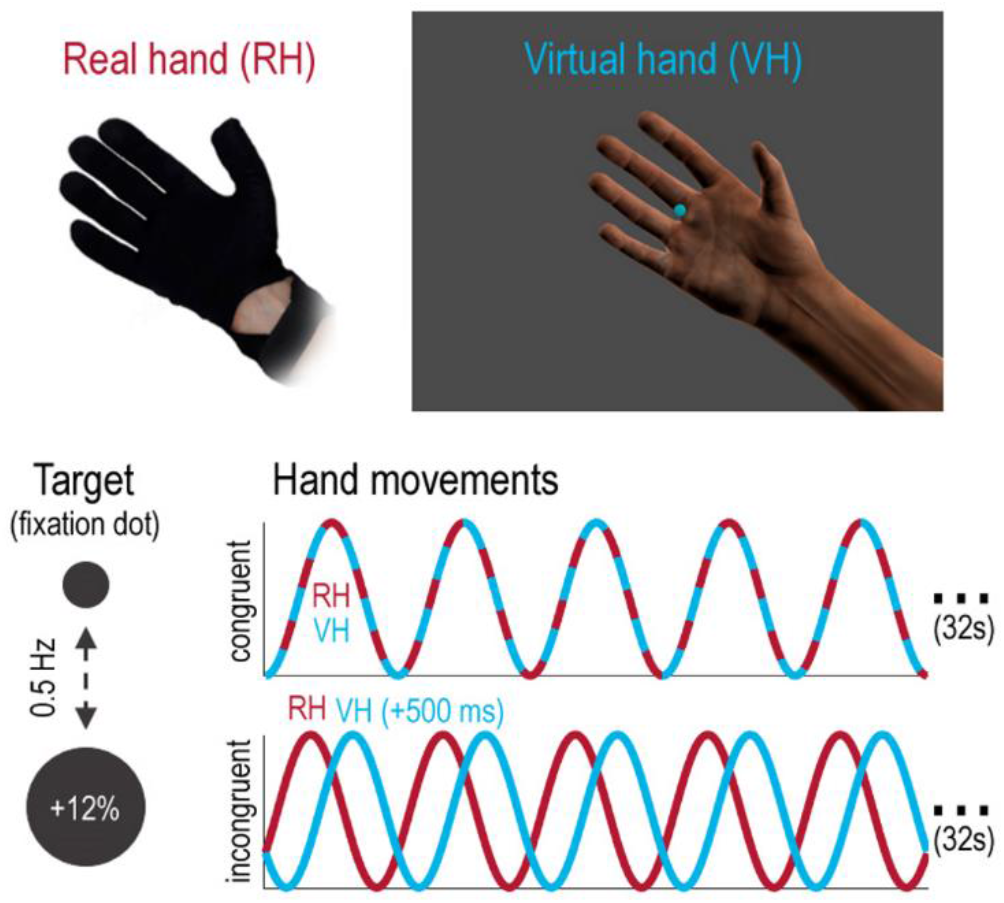
Phase matching task. Participants controlled a photorealistic virtual hand model (VH) with a data glove worn on their real hand (RH); the RH was occluded from view, while participants saw the VH at all times. Participants had to match the oscillatory phase of a virtual target (fixation dot, changing its size sinusoidally at 0.5 Hz) with right hand grasping movements (i.e., open at maximum target size, closed at minimum size). Thereby, participants were instructed to match the target’s oscillatory phase with the grasping movements of *either* the VH *or* the unseen RH. These instructions were intended to induce a specific task set, in which either visual or proprioceptive movement information was task relevant. In half of all trials, RH and VH moved congruently (‘congruent’), while in the other half of the trials (‘incongruent’), the movements of the VH were delayed (e.g., by 500 ms) with respect to the actually executed movements (RH); this introduced visuo-proprioceptive incongruence. Reprinted from Limanowski et al. (2020).

**Figure 2.**
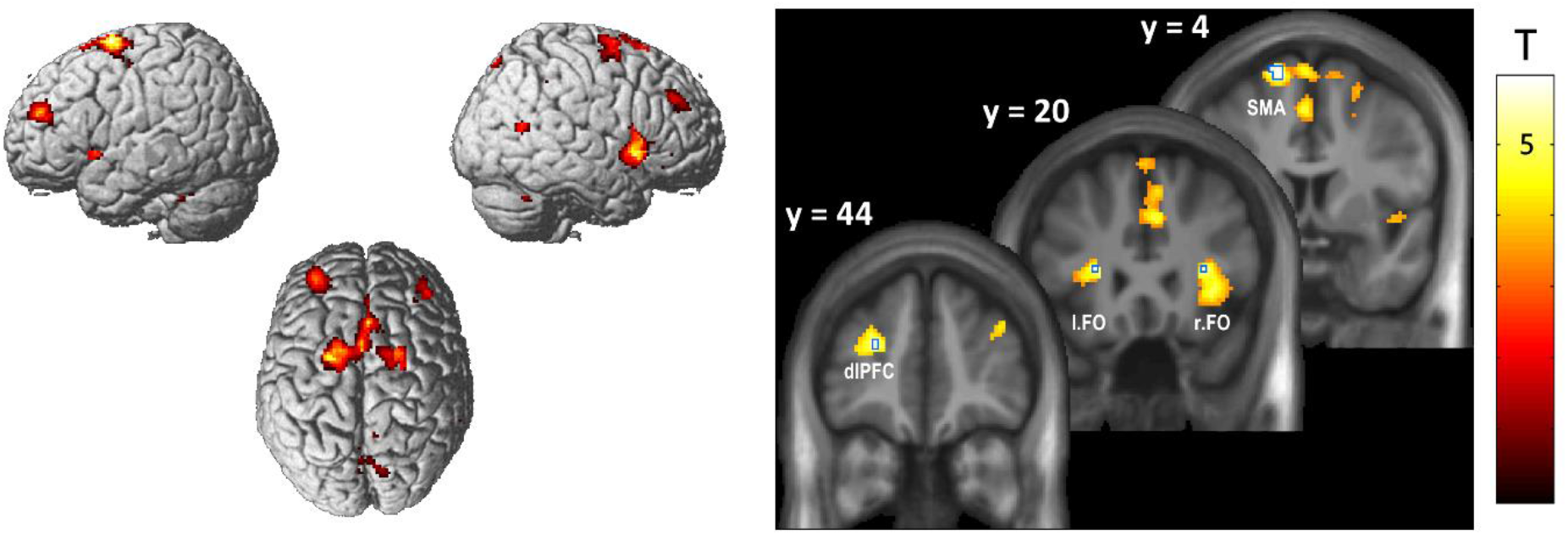
BOLD signal increases related to phase matching inaccuracy. The renders (left) and slice overlays (right) show brain areas in which hemodynamic activity was correlated with the relative inaccuracy of hand-target phase matching (displayed at *p* < 0.001, uncorrected). Significant activations (*p*_FWE_ < 0.05; voxels outlined in blue on the slice overlays) were located in the bilateral FO, the left SMA, and the left dlPFC.

On the group-level, the connectivity of the bilateral FO was evaluated using a paired t-test; i.e., a GLM including the PPI contrast images of the left and right FO of each participant, and another factor modelling the between-participant variance. We also tested whether the other two regions showing significant responses to phase matching inaccuracy (SMA and dlPFC) would exhibit connectivity changes, using a similar approach.

## Results

### Behavioural results

In both studies, participants were able to follow the task instructions; i.e., to keep the instructed modality’s (vision or proprioception) grasping movements aligned with the phase of the dot (see the original studies, Limanowski et al., 2020; Limanowski & Friston, 2020, for details and condition specific differences between task performance). In the MEG data, phase matching was on average significantly more variable in the RH compared to the VH conditions (*t*_(17)_ = 2.27, p < 0.05); but this was not the case in the fMRI data (*t*_(15)_ = 0.55, *n*.*s*.). On average, phase matching accuracy correlated weakly but significantly with movement amplitude (mean Pearson’s r = .27, *t*_(15)_ = 6.69, p < 0.001 for fMRI; mean *r* = .18, *t*_(17)_ = 5.08, *p* < 0.001 for MEG) but not significantly with the fMRI realignment parameters (all |*r*| < 0.02, *n*.*s*.).

### FMRI results

In our main fMRI analysis, we sought to identify brain regions in which neuronal activity correlated with phase matching (in)accuracy (Fig. 2). A significant (*p*_FWE_ < 0.05) main effect of inaccuracy was observed in the bilateral FO, the left SMA, and the left dlPFC (see Table 1 and Fig. 2). More liberal thresholds (*p* < 0.001, uncorrected) revealed further activation clusters in the right middle and superior frontal gyri, the precuneus, the medial cingulate cortex (MCC), the right middle temporal gyrus (MTG), and bilaterally in the cerebellum (cf. the render in Fig. 2). Conversely, a significant main effect of accuracy was found in the left pre- and postcentral gyrus, corresponding to the primary motor cortex (M1). No other comparisons (i.e., contrasting the effects of accuracy between task conditions, delay, or visual salience levels; see Methods) yielded significant effects. At uncorrected thresholds (*p* < 0.001), voxels in several brain areas showed a stronger correlation with task inaccuracy under the VH task than under the RH task; namely, in the MCC, the bilateral FO, the right MTG, the left cerebellum, the right dlPFC, and the bilateral posterior parietal cortex (PPC, peak within the intraparietal sulcus, IPS).

### MEG results

The MEG sensor space analysis revealed that phase matching inaccuracy was associated with significantly increased spectral power in the gamma frequency range over mid-frontal sensors (main effect; peak at 52 Hz, T = 5.31, *p*_FWE_ < 0.05; see Fig. 3A). These spectral effects were source-localized to the bilateral inferior frontal gyri, including the bilateral FO (Fig. 3B). No other spectral power comparisons yielded statistically significant results; but there was a statistical trend suggesting inaccuracy was associated with reduced alpha (8 Hz) power over posterior sensors (T = 4.61, *p*_FWE_ = 0.069).

**Figure 3.**
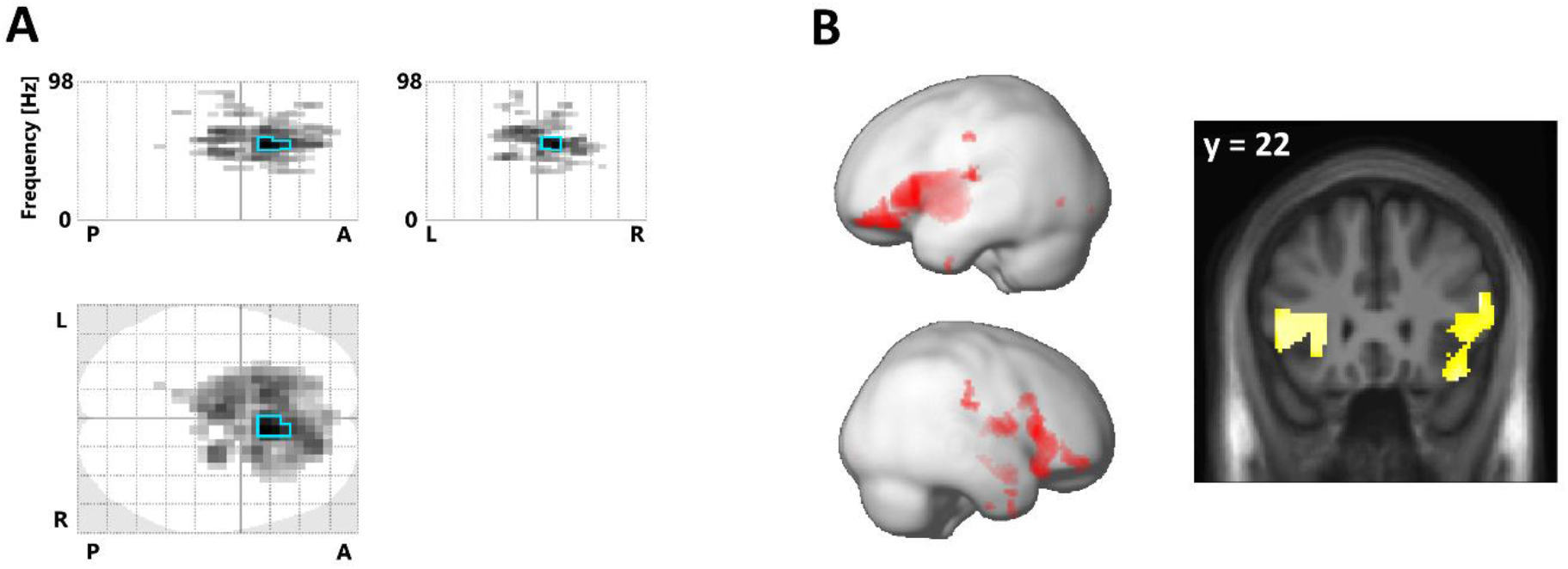
Spectral power increases related to phase matching inaccuracy. **A:** The ‘glass brain’ (maximum intensity) projections show the sensor level scalp-frequency maps of spectral power correlated with the relative inaccuracy of hand-target phase matching (the darkest voxels show the strongest effect along the respective projection; the maps are thresholded at *p* < 0.001, effects significant at *p*_FWE_ < 0.05 are outlined in blue; the top plots have one frequency dimension, 0 - 98 Hz, and one spatial dimension, P - A = posterior-anterior, L - R = left-right; the bottom plot has two spatial dimensions). **B:** Renders (left) and slice overlay (right) showing the corresponding source localization of the correlation to regions around the FO.

### Functional connectivity analysis

The above fMRI activations and source-localized MEG gamma power consistently suggested that periods of poor phase matching activated the bilateral FO, in line with previous literature that had established this region’s role in error processing and performance monitoring (see Introduction). However, we did not find any significant difference between conditions (i.e., between visual and proprioceptive task sets). Therefore, we next performed a connectivity (PPI) analysis on the fMRI data to explore whether task relevant brain areas would change their connectivity to the FO depending on the instructed task condition (VH or RH).

This analysis revealed a significantly increased coupling of several brain areas with the bilateral FO during the VH task > RH task, most strongly expressed in the right inferior parietal lobe (IPL, see Fig. 4A and Table 2). The increase in coupling with the right IPL was evident for both the left and right FO independently, as revealed by an additional ‘null’ conjunction analysis (Fig. 4B; i.e., a conjunction of voxels activated in the PPI with the left FO and PPI with the right FO, each thresholded at *p* < 0.001, uncorrected). Correspondingly, there were no significant differences in coupling between the left and right FO. A supplementary analysis testing for potential coupling differences with the FO during VH incong vs VH cong yielded no significant effects, either. There were no significant connectivity changes with the FO under the RH task > VH task. No significant changes in connectivity were observed in analogous analyses calculated for the SMA or the dlPFC, the other two brain regions showing significant effects in the main analysis (see above).

**Table 2.**
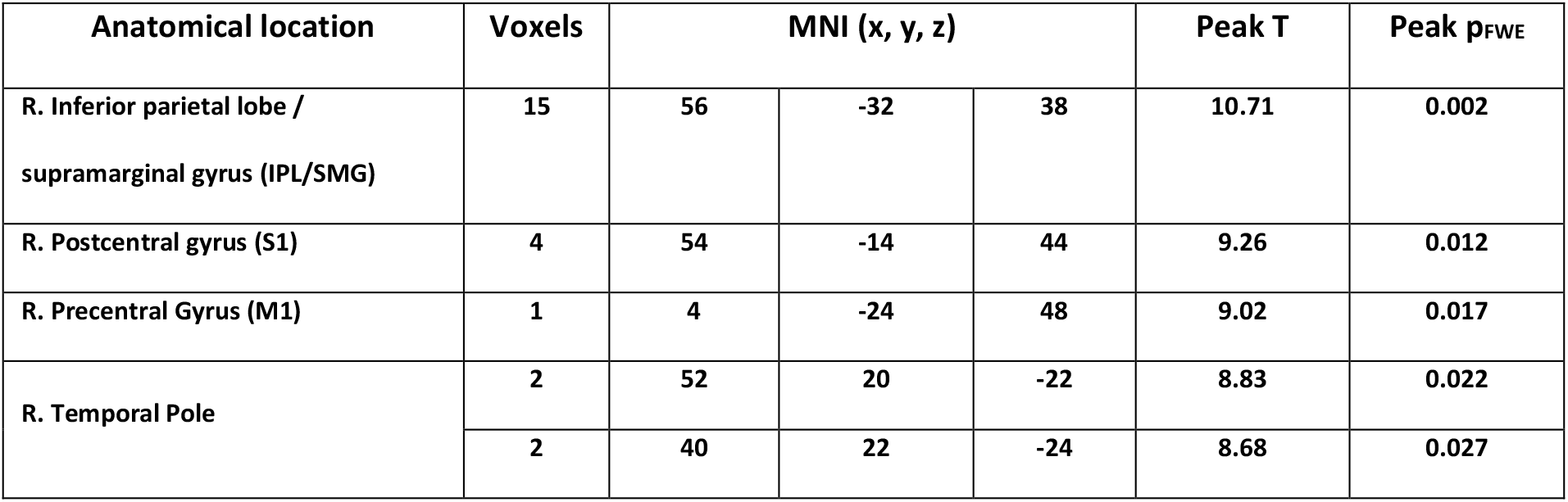
Brain areas showing significant (*p*_FWE_ < 0.05) coupling increases with the bilateral FO during the VH task > RH task.

**Figure 4.**
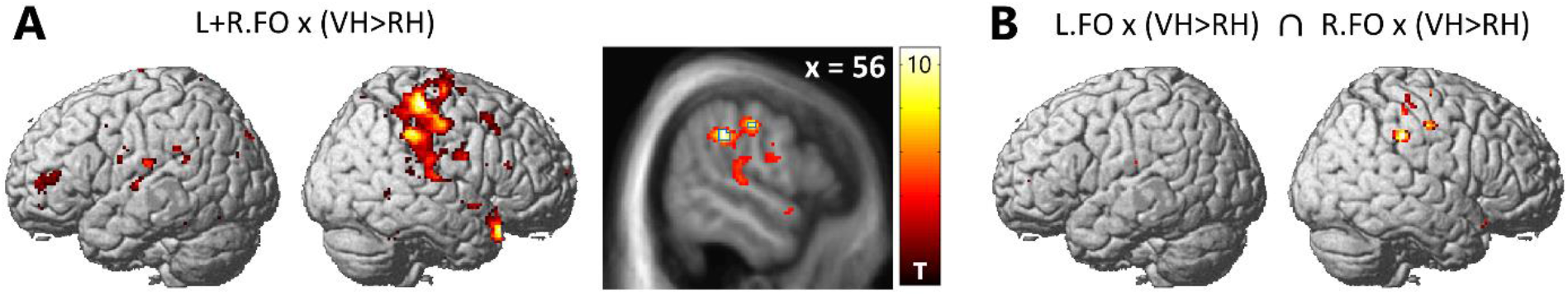
Task-dependent connectivity changes of the bilateral FO. **A:** Brain areas showing increased coupling with the bilateral FO during the VH task relative to the RH task (displayed at *p* < 0.001, uncorrected). The strongest effects were located in the right IPL (voxels significant at *p*_FWE_ < 0.05 are outlined in blue). **B:** A corresponding ‘null’ conjunction contrasts confirmed this increased task dependent coupling with the right IPL for the left and right FO independently (each PPI contrast thresholded at *p* < 0.001, uncorrected).

## Discussion

We used data from a virtual reality-based hand-target phase matching task to identify the hemodynamic and oscillatory correlates of performance (i.e., phase matching accuracy) monitoring under instructed task relevance of visual or proprioceptive hand position feedback. Our main result was a general, modality-independent response of the bilateral FO to poor phase matching accuracy, as evident from increased BOLD signal levels and increased gamma power. Furthermore, connectivity of the bilateral FO to the right PPC/IPL increased while participants executed the phase matching task with the visible virtual hand, compared to when they executed it with the real, unseen hand.

The observed general *BOLD signal increase* in the bilateral FO with task (phase matching) inaccuracy confirms observations of numerous previous studies, where BOLD signal in the FO increased in response to performance errors. For instance, the FO was activated by error trials vs correct trials in the Simon task (Danielmeier et al., 2011; Ham et al., 2013); in an antisaccade task (Klein et al., 2007); and in a flanker task (Eichele et al., 2008). Similarly, FO error-related BOLD signal increases were observed in visuomotor adaptation tasks (Grafton et al., 2008) and in response to tactile ‘oddball’ stimuli (Allen et al., 2016). Some studies found activation of the FO correlated positively with task performance (Bunge et al., 2015; Wager et al., 2005). This could, however, be explained with an general underlying function of FO activation in performance monitoring; acting not as an error signal per se, but as part of a mechanism to improve performance in response to errors (cf. Eichele et al., 2008). Thus, it has been proposed that neuronal activity in the FO may indicate the need for increased allocation of attentional resources to specific stimuli to achieve task-appropriate behaviour (Cieslik et al., 2015; Ham et al., 2013; Langner & Eickhoff, 2013; Uddin, 2021). Additionally, due to FO’s reciprocal connections to multiple sensory, limbic, and association areas (Sridharan et al., 2008), it may act as crucial ‘relay’ station for switching between different task relevant networks, e.g. switching from default network to executive control network (Klein et al., 2007; Menon & Uddin, 2010; Sridharan et al., 2008; Ullsperger et al., 2010).

The *spectral correlates* of task inaccuracy were expressed in the gamma frequency range; and, notably in agreement with the BOLD signal increases, they were source-localized to the bilateral FO (as part of larger sources in the IFG). A general correspondence and spatial co-localization of the BOLD signal and gamma power has been established in previous studies (Brovelli et al., 2005; Foucher et al., 2003; notably, including co-localization of responses in the insula, cf. Castelhano et al., 2014). Our findings align with previous studies that reported increases in intracranially recorded gamma activity in the FO following (stop-signal) task errors (Bastin et al., 2016). Moreover, gamma band activity per se is often interpreted as indicating enhanced processing of attended (e.g., task relevant) sensory information (Clayton et al., 2015; Hipp et al., 2012; Jensen et al., 2007; Siegel et al., 2012). In other studies, increased gamma power (over mid-frontal sensors) during response competition has been interpreted as indicating increased cognitive control (Grent-’t-Jong et al., 2013). A Granger causality analysis by (Chand & Dhamala, 2017) suggested that, during perceptual decision making, the FO may exert causal influence over fronto-parietal areas within the gamma band.

In sum, and in light of the above literature, our fMRI and MEG results suggest that the FO is involved in performance monitoring during goal-directed hand movements. Notably, while most of the above studies used trial-by-trial designs, our study featured continuous movements; thus, our results complement previous literature in showing that the FO shows similar responses in task settings requiring ‘on-line’ performance monitoring and adjustment during manual actions. Specifically, we propose that FO activation (expressed through BOLD signal and gamma power increase) may have indicated a reaction to task inaccuracy or error, and a corresponding need for behavioural adjustment. Tentatively, this interpretation is supported by the fact that posterior alpha power behaved opposite to gamma; i.e., it decreased with increasing inaccuracy (although this effect did not reach statistical significance, see Results). It is well established that posterior alpha inversely correlates with attention and task engagement (Bacigalupo & Luck, 2019; Sauseng et al., 2005; Thut, 2006; Yamagishi et al., 2003).

In addition to increased activation of the bilateral FO, our fMRI connectivity analysis revealed that these areas also increased their *functional coupling* with the right PPC (peak located in the IPL) during the VH task (phase matching with vision) compared with the RH task (phase matching with proprioception). An fMRI study by Higo et al. (2011) had used a task requiring attention to faces, houses, or body parts; and found that the FO increased its functional coupling with visual areas processing the respective task relevant stimulus category. In our case, however, the FO’s connectivity increase was not with primary and secondary visual cortices, which had shown task (attentional set) dependent activity increases in our previous studies (Limanowski et al., 2020; Limanowski & Friston, 2020). Instead, FO coupling increased with the IPL of the right PPC; an area that is involved in more high-level processes including multisensory and sensorimotor integration, and visuo-spatial attention (Andersen & Buneo, 2002; Malhotra et al., 2009; Wolpert et al., 1998).

We propose that this result is related to the fact that visual hand movements were task relevant in the VH task, but had to be ignored in the RH task (where phase matching was done with proprioception). Thus, visual feedback was essential for correcting phase matching error in the VH task, but irrelevant in the RH task. Notably, FO connectivity was not significantly different during periods of visuo-proprioceptive incongruence; neither did we find significant differences between congruent and incongruent conditions in the main fMRI GLM analysis. This suggests that the observed VH > RH task dependent connectivity difference was related to the task-relevant modality being vision > proprioception per se, rather than to (in)congruence between vision and proprioception. This interpretation fits with previous work showing that the right PPC, specifically areas in the right IPL, are critical hubs for executing and correcting visually guided arm movements (Culham et al., 2003; Culham & Valyear, 2006; Desmurget et al., 1999; Lane et al., 2011; Ogawa et al., 2007; Wenderoth et al., 2004).

Potentially, this effect might have been enhanced by intrinsic differences between the modalities in relation to error detection; i.e., it might have been easier for participants to notice a phase matching error when focusing on visual action feedback (VH task) than when focusing on proprioception (RH task). This could have been due to visual body position estimates being intrinsically less variable than proprioceptive ones (cf. van Beers et al., 1999); and because visual body position was easier to compare to the visually presented target (however, the target quantity was not visuo-spatial but abstract; i.e., the oscillatory growing-and-shrinking phase of the fixation dot). When we lowered the statistical threshold of the main GLM analysis to *p* < 0.005, uncorrected, the bilateral FO (the PPI seed regions) and the right PPC (the PPI target region) showed a stronger correlation with task inaccuracy under the VH task compared with the RH task. Although this was a weak effect, it could mean that overall, errors were more easily processed (in those areas) in the VH task. Interestingly, participants were overall worse in the RH than in the VH task, and performance varied more strongly in the RH task; this could support the interpretation that proprioceptive performance monitoring was less efficient than when vision was used. This may also fit with previous reports of increased BOLD signal in the FO for error trials of which participants were aware, than unaware errors (Harsay et al., 2018; Klein et al., 2007). Specifically, Harsay et al. (2018) also observed increased functional connectivity of the FO to the PPC (bilaterally, in addition to the bilateral S1) during aware > unaware errors. In sum, we suggest that the increased connectivity between the FO and the right PPC during the VH > RH task indicates the FO signalling an increased need for control and attentional and/or behavioural adjustment (following poor performance) to visuomotor regions in the right PPC; which could also be related to how easily those performance deficits could be detected.

The above speculations could also explain why we did not observe any connectivity increases of the FO during the RH > VH task. Accordingly, this could be because participants were less aware of their phase matching (in)accuracy when performing the task with the unseen real hand. Future work should evaluate this possibility with specific task designs.

Besides activations in the bilateral FO, we also found BOLD signal increases to poor phase matching in the SMA and the dlPFC (at the junction of middle frontal gyrus and frontal pole). Both areas have been strongly implied in performance monitoring in other contexts (Ullsperger et al., 2014). The SMA has been shown to respond to unexpected stimuli, e.g. surprising action outcomes (Krakauer et al., 2004; Sakai et al., 1999; Scangos et al., 2013; Ullsperger et al., 2010). BOLD signal in the SMA has previously been reported to correlate with positional error in a visuomotor learning task (Grafton et al., 2008) and in a continuous hand-target tracking (Limanowski et al., 2017). In line with the interpretation provided in these studies, the SMA activation we observed may indicate an updating of movement plans in response to poor detected phase matching. Similarly, the lateral PFC is considered a crucial part of sensorimotor hierarchy (Benchenane et al., 2011; Sokhadze et al., 2012), and is thought to contribute to performance monitoring and error detection; e.g., by preparing attentional task sets and comparing behavioural output against them (Cieslik et al., 2015; Danielmeier et al., 2011; Smith et al., 2019; Ullsperger & von Cramon, 2004). In our experiment, the dlPFC activation could imply similar underlying ‘high level’ functions.

Conversely, we observed that BOLD signal in the contralateral M1 correlated positively with task accuracy. This effect could be related to the fact that higher task accuracy coincided with more pronounced hand movements; however, we had included movement amplitude as a regressor of no interest in our first-level GLMs, which should have largely accounted for this potential bias. Alternatively, this observation also aligns with the M1’s known role in motor learning (Hardwick et al., 2013; Panico et al., 2021; Spampinato & Celnik, 2017); with previous findings that M1 activity correlated with visuomotor adaptation performance (Della-Maggiore & McIntosh, 2005) or with visuomotor target tracking performance (Ogawa et al., 2006); and with the fact that a perturbance of the M1 via transcranial magnetic stimulation (TMS) resulted in reduced sensorimotor adaption (Orban de Xivry et al., 2011).

Finally, it should be noted that our results should be compared to previous studies with some caution, since our task was designed around continuous movements; therefore, we could not isolate specific time points—and neuronal correlates—that would clearly correspond to specific cognitive or motor processes like e.g. error detection or correction. Future trial-by-trial task designs should therefore try to validate our interpretation.

In conclusion, our results suggest a critical role for the bilateral FO in performance monitoring during manual action; and that following errors in visually guided manual action specifically, the FO may signal an increased need for control to visuomotor regions in the right PPC.

## Acknowledgments

Funded by the German Research Foundation (DFG, Deutsche Forschungsgemeinschaft) as part of Germany’s Excellence Strategy – EXC 2050/1 – Project ID 390696704 – Cluster of Excellence “Centre for Tactile Internet with Human-in-the-Loop” (CeTI) of Technische Universität Dresden. JL is funded by a Freigeist Fellowship of the VolkswagenStiftung (AZ 97-932). Figure 1 reprinted from: Limanowski *et al*., Neuroimage, 222, p. 117267, Copyright Elsevier (2020) under the terms of the Creative Commons CC-BY license.

